# Template-assisted fabrication of moon-shaped channels for protein breakthrough analysis

**DOI:** 10.1101/2023.09.14.557754

**Authors:** Raghu K. Moorthy, Serena D’Souza, P. Sunthar, Santosh B. Noronha

## Abstract

Cylindrical column with packed stationary phase is the workhorse of liquid chromatography systems. These stationary phases are commonly classified on the basis of different form factors namely, beads and monoliths for protein chromatography. Monolithic rods are one of the important geometries derived from polymers through complex polymerization schemes with additional requirements such as cross-linkers and specific reaction conditions. To address these practical difficulties and enable ease of fabrication at laboratory scale, acrylic copolymers are hypothesized to perform as a monolithic stationary phase suitable for protein chromatography. The present work proposes a rapid fabrication technique to obtain monolithic rods that could be reconditioned without any of the above additional steps. It is characterized with monolith diameter that could be controlled using acrylic copolymer concentration. Formation of the copolymeric stationary phase inside microchannel led to annular geometry and in turn, demonstrated fabrication of moon-shaped channels (MSCs) for the first time in literature. An online monitoring system facilitated tracer breakthrough analysis with MSCs to report sharp peak front and an estimate of channel void volume. Breakthrough curves with single protein validated the selection of blue dextran as tracer and indicated retention of proteins due to electrostatic interactions on the functional copolymer surface.

## 1. Introduction

Stationary phase in liquid chromatography columns is an integral part of chromatography-based separation techniques to resolve analytes of both small and high molecular weight. Different working principles like ion exchange, affinity or both modes are leveraged to elute analytes of interest preferentially using suitable functional materials [1]. Examples include polymer resins (acrylamide, methacrylate-derived polymers), graphite carbon, carbon siloxanes and silica particles [2]. The stationary phase is packed into conventional chromatography column using specialized equipment at high pressure and temperature in centralized, commercial facilities ([1]; [3]). In protein chromatography, different types of stationary phase mainly consist of spherical beads and monoliths derived out of polymers and its (chemically) functional derivatives ([2]; [4]; [5]). To reduce pressure drop across protein chromatography columns in particular, packed monolith as continuous rod is fabricated inside tubular columns to represent the stationary phase ([4]; [6]; [7]). Capillary dimensions (less than 1000 *μ*m diameter) are preferred as tubular columns to develop different protein separation technology platforms as first reported in the pioneering work by Svec and Fretchet in the 1990s. Over the past three generations, these monoliths have evolved into various categories on the basis of different tubular channel configurations and type of monolith, typically using modified silica or copolymers ([6]; [8]; [9]; [10]; [11]) (Table 1 and Figure 1). In all of the above case studies, formation of stationary phase as monoliths required high temperature (between 50 to 600*°*C) and reactive polymerization conditions such as addition of chemical cross-linkers.

**Table 1:** Case studies for fabrication of monoliths used in liquid chromatography.

| Monolithic material | Polymerization required* | Reaction temperature during monolith formation (°C) | Reference |
| --- | --- | --- | --- |
| Poly(glycidyl methacrylate-co-ethylene dimethacrylate) | Yes | 70 | [4] |
| Octadecyldimethyl-(N,N-diethylamino) silane (ODS)-modified silica | Yes | 50 to 600 | [19] |
| Silanized copolymer of lauryl methacrylate and 1,6-hexanediol | Yes | 50 to 120 | [20] |
| Octadecylsilylated (ODS)-modified silica | Yes | 60 or 87 | [21] |
| Poly(styrene-co-divinylbenzene) | Yes | 70 | [22] |
| Poly(butyl methacrylate-co-methacrylic acid-co-ethylene glycol dimethacrylate) | Yes | 53 | [23] |
| Poly(2-vinylnaphthalene-co-divinylbenzene) | Yes | 65 | [24] |
| Acrylic copolymer (ACP) | No | 25 to 30 | Present work |

\*Polymerization using monomers is required as one of the steps toward monolith formation.
ACP - Poly(methyl methacrylate-co-methacrylic acid)

**Figure 1:**
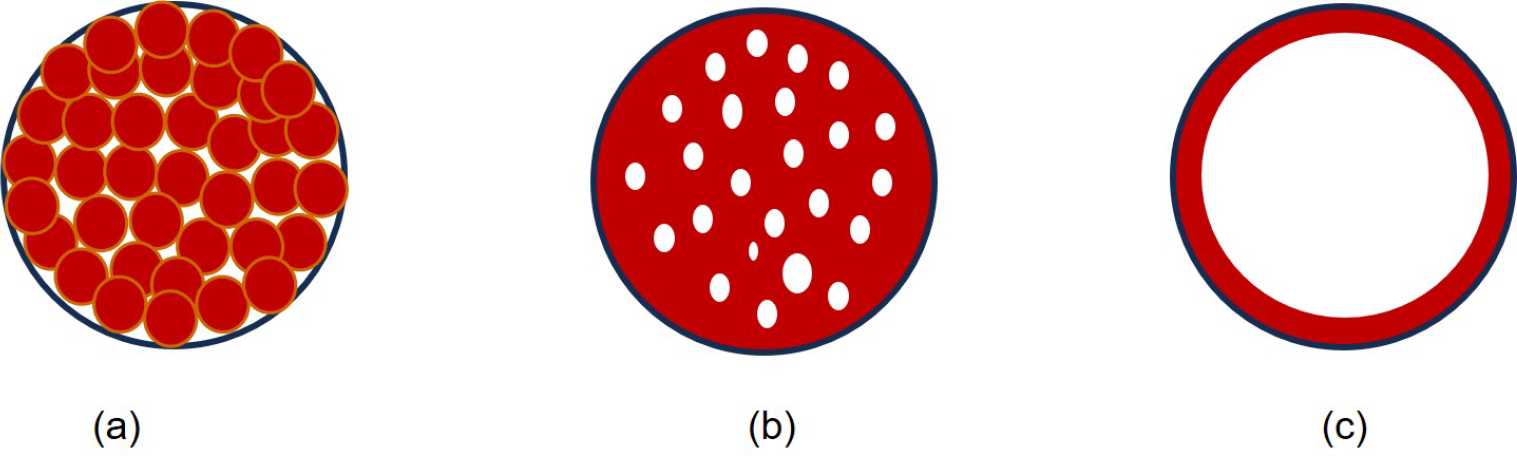
Schematic diagram of different types of protein chromatography columns: (a) Packing/bead-based (b) monolith (c) wall coated.

In liquid chromatography, annular geometry has been utilized to improve dispersion characteristics. It provides different angular distances when combined with rotating packed bed configuration [12]. The inner core in this arrangement is usually the stationary phase (porous or non-porous) through which analytes from bulk liquid flow through at different retention times, depending on both void volume and electrostatic interactions with the functional ligands. Recently, there is theoretical evidence that stationary phase (porous or non-porous) as fixed inner core in such annular geometry reduces dispersion [13]. Cylindrical columns with a perfect cylindrical fixed core (porous or nonporous) has been a manufacturing challenge and therefore, alternative geometry for inner core along with different aspect ratio have been studied by various research groups over the past five decades ([14]; [15]). Moon-shaped channels based on different arc radius of the inner core have been predicted to provide residence time distributions with reduced dispersion [15] and resultant breakthrough curves for protein chromatography applications. Online monitoring of proteins eluting out of chromatography system is then required to identify these intrinsic flow characteristics using a concentration step input at column inlet (or continuous supply of analyte as feed) [16]. For mass diffusion-controlled processes as in protein chromatography, residence time distribution (RTD) provides useful insights on the effect of dispersion at a given inlet flow rate. Fraction of molecules that spend a certain time inside the channels (denoted as ‘E’) could be estimated and represented in the form of a distribution chart or E-curve [17] after mathematical transformation of breakthrough curves (or F-curves) using first order differential derivatives [18].

In this work, formation of copolymeric stationary phase as monolithic rod inside microchannel using template-based micromolding as a rapid fabrication technique is proposed. Ease of fabrication by use of ambient conditions and no additional requirements as in conventional packing methods are hypothesized to obtain the monolith at a laboratory scale.

## 2. Materials and methods

### 2.1. Chemicals and reagents

Polydimethyldisiloxane (PDMS) elastomer kit (Sylgard 184) was supplied from Dow Corning Incorporated, USA. Monosodium dihydrogen phosphate (Cat No. 59443) and disodium hydrogen phosphate (Cat. No. 21669) (all an-hydrous) (Sisco Research Laboratories, India) for phosphate buffer preparation were purchased. These chemicals were used as received without further modification. Poly(methyl methacrylate-co-methacrylic acid) (85:15 monomer ratio) (Cat. No. 376914) (hereafter, referred to as acrylic copolymer (ACP)) (Sigma, USA) was purchased. Model protein (supplied in lyophilized powder) used in breakthrough analysis was human hemoglobin (Hb) (Cat. No. H7379) and human hemoglobin A_0_ (HbA_0_) (with 5% (w/w) methemoglobin) (Cat. No. H0267) (all protein samples from Sigma, USA). Tetrahydrofuran (THF) was the organic solvent used for ACP dissolution. 5 M sodium hydroxide (NaOH) and 1 M hydrochloric acid (HCl) solutions were prepared and used in channel fabrication procedure [25] and for adjustment of pH during preparation of phosphate buffers respectively. The prepared solutions were filtered and sonicated before use. All analytical grade chemicals or reagents unless stated otherwise were purchased from Merck Laboratories, India. Deionized (DI) water (Millipore, USA) has been utilized throughout the experiments.

### 2.2 Preparation of moon-shaped channels (MSCs)

#### 2.2.1. Template-assisted micromolding

Template-assisted micromolding method has been performed to obtain microchannels embedded in a microchip, devoid of any bonding requirements. Acrylic sheets were cut out using laser ablation technique (Suresh Indu Lasers (SIL), India) to fabricate a rectangular enclosure. Holes of 1 mm were drilled through 4 mm thick acrylic sheet to place metallic needles (see Figure 2 below). Stainless steel needles (Becton Dickinson (BD), Singapore) were used to tightly stretch out nylon thread along the length of the enclosure. Here, nylon thread (Golex, India) represents the template for a microchannel. Prepare degassed PDMS solution (max. vacuum 600 mm Hg, Tarsons Rocker 300, India) using silicone elastomer and curing agent mixed in the ratio varying between 9.5:1 and 10:1. PDMS was poured over the thread to fabricate a microchip of about 4 mm thickness. The setup was placed on a hot plate (C-MAG HS 7 Digital, IKA, Germany) at a temperature of 75 ^*°*^ C for nearly 2 hours. On cooling (about 30 minutes), the nylon thread was smoothly pulled out and PDMS microchip was then carefully peeled off on cooling to obtain a solidified, hollow cylindrical microchannel. Inlet and outlet reservoirs of 1.5 mm diameter were punched in-line to the ends of microchannel. Microfluidic connectors were utilized to connect silicone tubing with the microchannel for liquid flow in further use.

**Figure 2:**
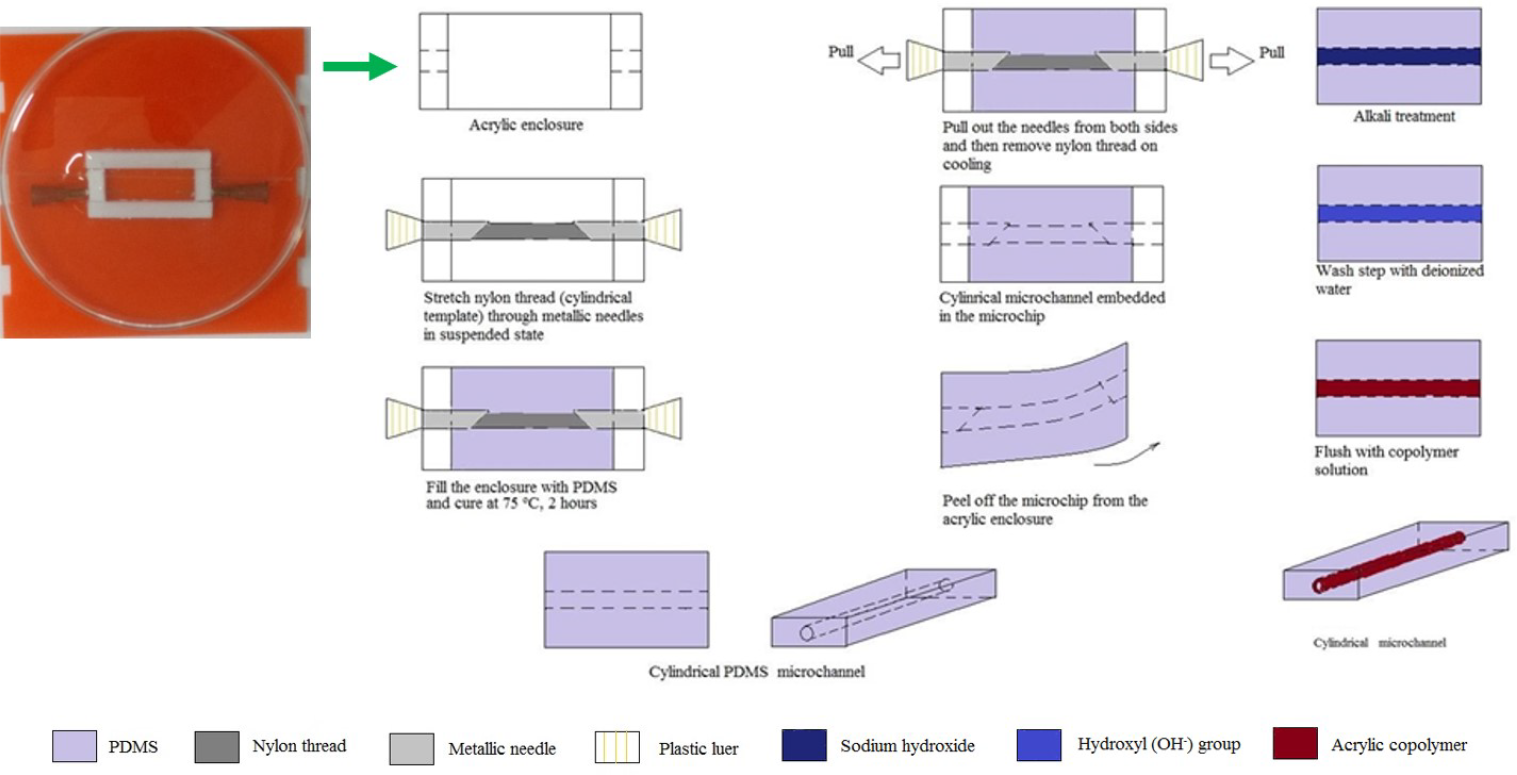
Template-assisted micromolding: Acrylic template enclosure fabricated for curing channel wall substrate.

#### 2.2.2. Monolith formation

A microfabrication method was developed to obtain PDMS microchannels with a porous polymer cylindrical rod as monolith. Extending along the microchannel length, the rod was to be anchored at base (or bottom side) of the microchannel. Prerequisites for this process include preparation of (a) different concentrations (in %(w/v)) of acrylic copolymer solution (ca. 0.5 g poly(methyl methacrylate-co-methacrylic acid) dissolved in 5 ml tetrahydrofuran (THF) for 10%(w/v)) [26] and leave it undisturbed after initial mixing ∼ 5 to 6 hours for complete dissolution and (b) PDMS microchannel (170 *μ*m) using the template-based micromolding method, as mentioned in the earlier section. 5 M sodium hydroxide solution was then passed through the microchannel at an inlet flow rate of 10 *μ*g ml^−1^ for about 15 to 20 minutes using syringe pump (NE-300, New Era Pump Systems, USA). The channel was washed with deionized water at 10 *μ*l min^−1^ for 5 minutes (add 10 minutes for water priming in the inlet tube). Connect the inlet of microchannel directly to the syringe loaded with 10%(w/v) acrylic copolymer using a (1/16)” microfluidic connector (WW15954-91, Cole-Parmer, USA). Flush the channel using the above solution at 100 *μ*l min^−1^ for 30 s or until the first drop of solvent arrives at the outlet, whichever is later. Repeat this step for one more time immediately and leave it for solvent drying overnight (at least 24 hours) before further use.

### 2.3 Scanning electron microscopy (SEM)

The microchannels were prepared for imaging through a 45 ° or 90 ° section cut straight across the microchannel cross-section (see Figure 3). Different regions of interest along the microchannel length were considered to view surface morphology of the monolith inside the microchannel. Surface imaging of the untreated and treated PDMS microchannels were performed using field emission scannig electron microscope (FESEM, JEOL JSM-7600F, Japan) at accelerating voltage of 5 kV. The polymer surface of interest was coated with a thin film of platinum to negate surface charging (film thickness of less than 10 nm). The images have been captured at a working distance ranging between 5 to 26 mm away from the sample surface.

**Figure 3:**
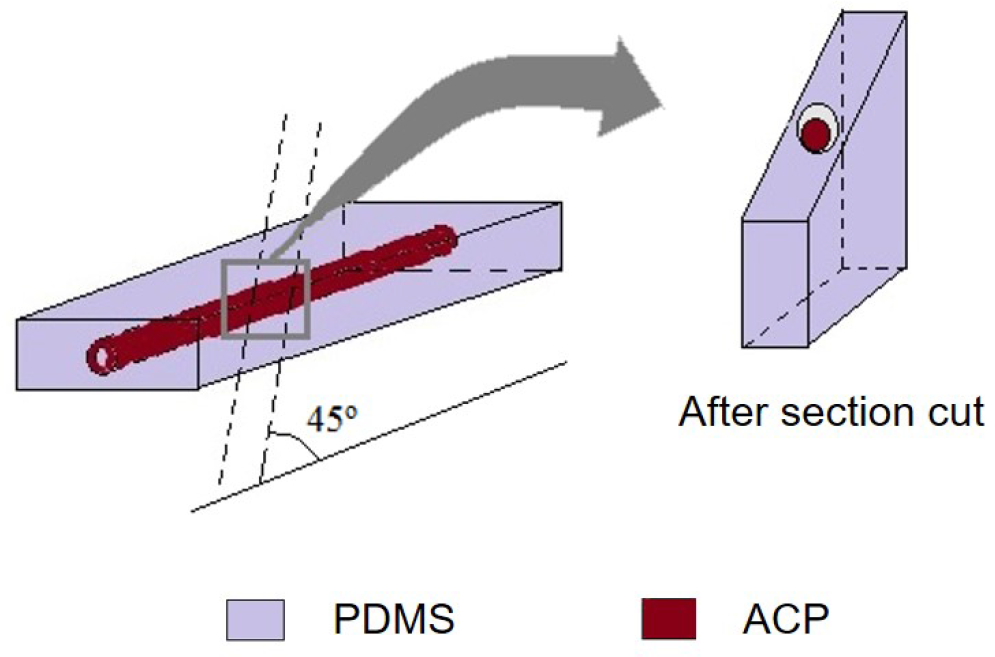
Sample preparation for SEM analysis of cylindrical microchannels after monolith formation using acrylic copolymer (ACP).

### 2.4. Image analysis

Calibrated scales for each electron micrograph were set in the order of tens of micrometers at low magnification. The channel diameter (D_c_) and monolith diameter (D_m_) were measured to estimate an annular diameter (D_c_ − D_m_) and corresponding geometrical area of cross-section (ImageJ, NIH, USA). Thresholds were set for all SEM images and porosity of monolith estimated. For a given scanned image, total area was set to 100% and a relative area fraction (in per cent) was measured using three different internal regions of interest. Additionally, SEM images of cross-section of microchannels using 10% acrylic copolymer were verified for anchorage to channel walls at 700X magnification and nearly no flow area was to be established. In case of visual inspection of the channels before and after monolith formation, a low magnication of 10X using simple microscope (Leica DM500) was utilized.

### 2.5. Online monitoring system

The main components of online monitoring system include: (a) syringe pump (b) silicone tubing (1 mm diameter) (c) stainless steel capillary connector (0.4 mm diameter) (d) detector flow cell (10mm path length, 17 *μ*l cell volume) (e) light source as deuterium or tungsten lamp for a UV-Vis wavelength range (195 to 650 nm). Moon-shaped channels (MSCs) or channels treated without ACP or untreated channels (as negative controls) were plugged in between the pair of silicone tubings and connectors to complete the system connections (Figure 4).

**Figure 4:**
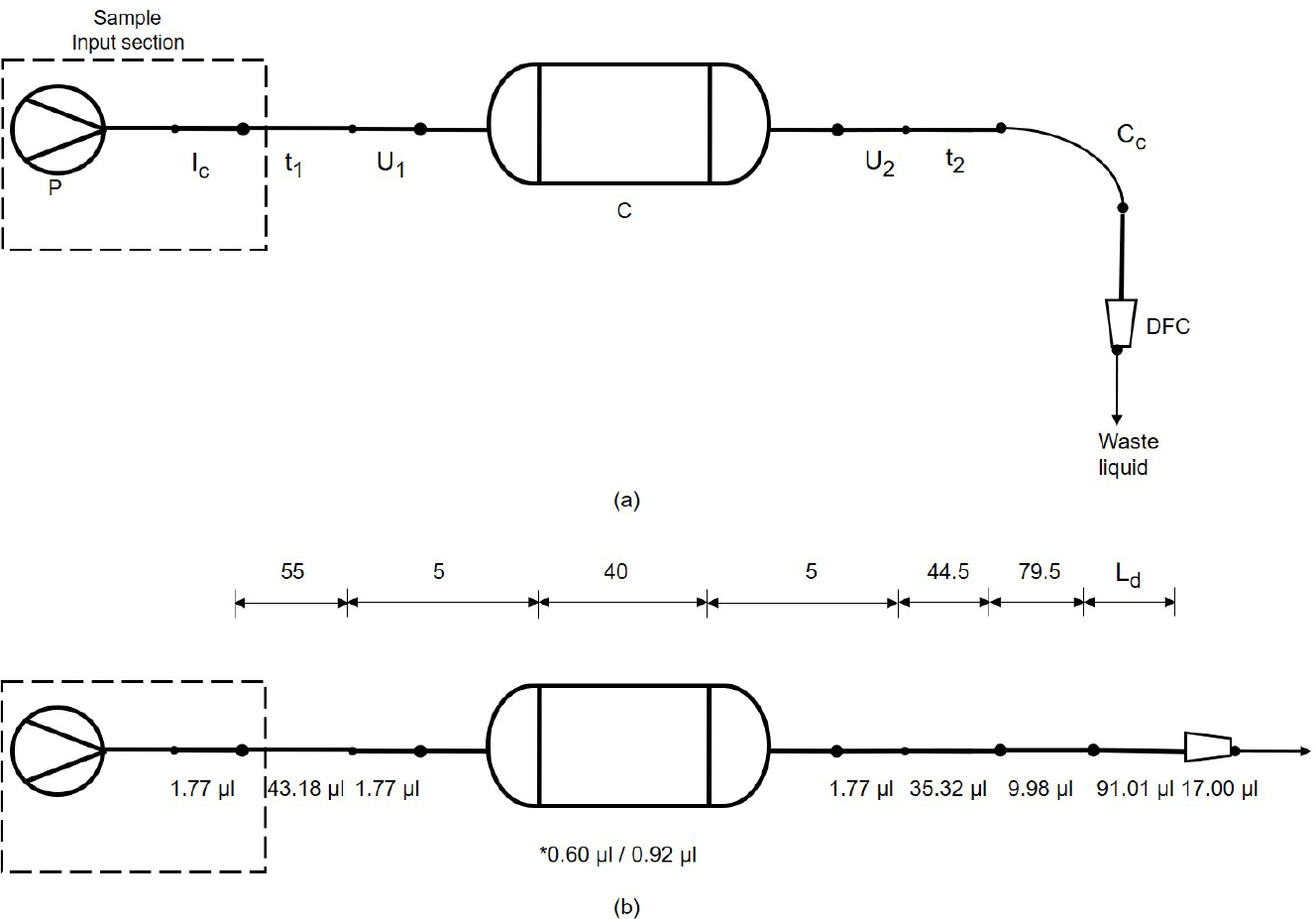
Schematic representation of current microfluidic experimental setup: (a) flow scheme with different components, P syringe pump, I_C_ inlet connector, C column (untreated microchannel, treated microchannel devoid of ACP or *moon-shaped channel (MSC)), U_1_ - union 1, t_1_ - inlet tubing, U_2_ - union 2, t_2_ - outlet tubing, C_C_ - capillary connector (upto detector inlet), L_d_ - unknown flow length inside detector, DFC - detector flow cell (b) diagram showing different dimensions (in millimeters) and volume contribution for each component. Note: dimensions are not to scale.

**Figure 5:**
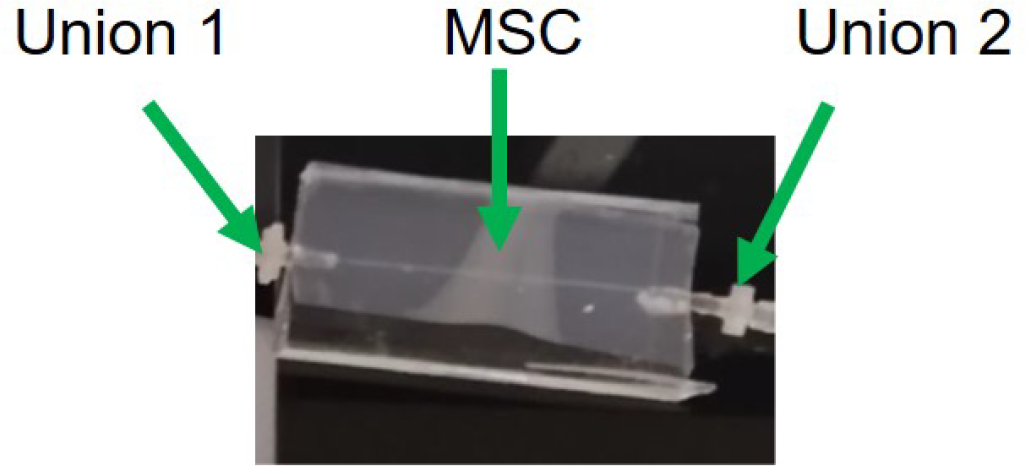
Photograph of freshly prepared moon-shaped channel (MSC) embedded in a PDMS microchip with inlet and outlet connectors.

The absorbance data was recorded at a sampling rate of 1 data point per minute of analysis time. The detector accumulation time for the data was set at 800 ms (less than 1 second). In particular, tracer (blue dextran) and proteins as analytes were monitored at 615 nm and 280 nm respectively.

### 2.6. System volume

Apart from moon-shaped channel (MSC) as mentioned earlier, the system volume consists of the following flow sections: inlet tubing length = 55 mm, outlet tubing length (from outlet of MSC upto online detector) = 124 mm. The separation system assembled for further experimentation has been shown in Figure 4.

### 2.7. Breakthrough analysis

#### 2.7.1 Tracer studies

MSCs of nearly 170 *μ*m diameter and 40 mm length were fabricated using 10% acrylic copolymer concentration (in (w/v)%) and connected within the online monitoring system as described earlier (Section 2.5). Inlet and outlet tubings and connectors (Figure 4) were subjected to phosphate buffers as solvent used in these stuides, namely, buffer MB (20 mM phosphate buffer, pH 6.4) and buffer BD (Blue dextran in buffer MB). As part of initial priming, buffer MB was flown inside the system till the first liquid drop comes out at the outlet tubing. The entire annular space available inside the column was filled with buffer MB after following the above steps. Continuous supply of blue dextran solution (buffer BD containing 2 mg ml^−1^ blue dextran) as step input was loaded directly at the inlet tubing after priming the system with buffer MB. The above continuous flow of buffer solution inside the column was carried out at an inlet flow rate of 5 *μ*l min^−1^ until blue dextran concentration reached a saturation level. The liquid fractions were analyzed in-line using online monitoring of abosrbance of blue dextran at 615 nm (Intelligent UV-Vis Multiwavelength detector, Part number: 0302-0546A, Jasco MD-2010 Plus, Japan) after standard calibration. For control purposes, above experimental system without connecting the column was assembled and rest of the procedure was carried out.

#### 2.7.2. Protein studies

Inlet or outlet tubings, MSC and connectors as part of the online monitoring system were arranged as described earlier (Section 2.7.1). The system tubing was subject to wash in multiple steps with deionized water. Protein solution (buffer BP containing model protein (human hemoglobin or Hb) of nearly 1000 *μ*g ml^−1^ feed concentration in 20 mM phosphate buffer at pH 6.4 (buffer MB)) was loaded directly at the inlet tubing after priming the column with 20 mM phosphate buffer at alkaline pH 9.3 (buffer AB). Buffer AB was flown through till the first liquid drop came out at the outlet tubing. Absorbance of liquid eluting at channel outlet was monitored using an inline flow cell arrangement (Intelligent UV-Vis Multiwavelength detector, Part number: 0302-0546A, Jasco MD-2010 Plus, Japan). Thus, the entire available annular space inside the column was filled with presaturation buffer (buffer AB) until baseline signal at 280 nm was stabilized. Continuous supply of buffer BP (as step input) inside MSC was carried out at an inlet flow rate of 5 *μ*l min^−1^ until protein concentration reached a saturation level (final) after an initial start from breakthrough (at around 10% tail height) point.

### 2.8. Elution of total protein inside MSCs

The system tubings, MSC and connectors as part of the microfluidic experimental setup were arranged as earlier (Section 2.7). Protein solution (buffer BP containing model protein (human hemoglobin A_0_ or HbA_0_) of nearly 1.7 *μ*g total protein in 20 mM phosphate buffer at pH 6.4 (buffer MB)) was injected as pulse input (analyte volume 1 *μ*l) using a glass syringe (50 *μ*l glass syringe 705N, Part number: 80565/00, Hamilton, USA) at the inlet tubing after priming moon-shaped channel (MSC) with 20 mM phosphate buffer as presaturation buffer of pH 2.5 (buffer PB). Buffer PB was flown through till the first liquid drop came out at the outlet tubing and until baseline signal at 280 nm was stabilized. It was followed by the pulse input of buffer BP within a narrow time span of less than 30 seconds. A continuous step input of buffer PB was carried out at an inlet flow rate of 5 *μ*l min^−1^ for nearly three hours from start of the syringe pump. Absorbance of liquid eluting at channel outlet was monitored using an inline flow cell arrangement. Continuous supply of mobile phase buffer (buffer MB) (as step input) inside MSC was carried out at an inlet flow rate of 5 *μ*l min^−1^ immediately after from two hours of analysis time until nearly complete elution of total protein was obtained.

### 2.9. Residence time distribution for system and MSCs

In residence time distribution (RTD) studies, an inert material (tracer) is considered as analyte for obtaining concentration-time data which is then, transformed into E-curve using the following equations:

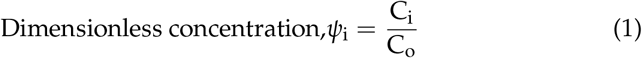

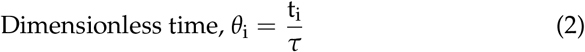

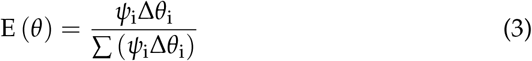

where;

C_i_ = Analyte concentration (after step input feed) at time t_i_

C_o_ C_i_ = Analyte concentration (after pulse input feed) at time t_i_

C_o_ = Analyte feed concentration (*μ*g ml^−1^)

t_i_ = Individual analysis time (min)

*τ* = Mean residence time inside chromatography system 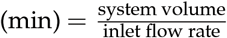

Dimensionless analyte concentration at outlet is then denoted as ‘F’ and could be transformed back into E-curve using the following equations:

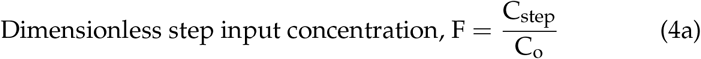

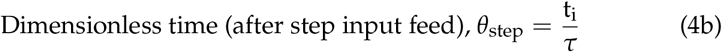

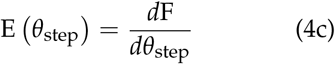

where;

C_step_ = Analyte concentration (after step input feed) at time t_i_

For ideal laminar flow, tubular column with maximum superficial flow velocity at center of tube, RTD is similar to a combination of impulse with a long tail. It implies highest fraction of molecules elute out within a short residence time if convective forces are dominant over diffusive forces (including axial dispersion characteristics).

## 3. Results and Discussion

### 3.1. Surface morphology inside moon-shaped channels (MSCs)

The functional surface (or ACP) formed as monolithic stationary phase inside cylindrical columns has been defined as a porous cylindrical rod extending along the channel length. The micropores were to be distributed over its surface as similar to the monolithic structures (Table 1). In the present work, a narrow annular segment was formed with alkali-treated PDMS surface on one side ([25]; [27]) and stationary phase surface anchored on the other side. Through a series of scanning electron microscopy (SEM) images, observations have been recorded for different acrylic copolymer (ACP) concentrations (5, 10, 15% (w/v)), the cross-section area occupied by the stationary phase increased with increase in ACP concentration (Figure 6). To the best of our knowledge, fabrication of these columns as moon-shaped channels (MSCs) has been demonstrated for the first time in literature. An important feature of MSC was that with only a change in monoliith diameter (or geometric arc radius) using different ACP concentration, a family of MSCs could be fabricated. Since amount of ACP increased with its concentration, larger area of cross-section inside the channel was covered by formation of stationary phase as cylindrical, monolithic rod after near complete evaporation of the organic solvent. Using an online moniitoring system, these MSCs were demonstrated to be low pressure columns with low porosity inside stationary phase (less than 10% of available surface area) and suitable for lower operating flow rates of less than 10 *μ*l min^−1^. By geometrical calculations, for MSCs of 40 mm length and annular diameter (*D*_c_ − *D*_m_) of around 70 *μ*m in the case of 10% (w/v) ACP concentration, where *D*_c_ ∼ 170 ± 2 *μ*m (measured from 100 different microchannels (Leica DM500 and LAS 3.4 Application software)) and *D*_m_ ∼ 100 *μ*m (measured from SEM images (ImageJ, NIH, USA)); channel void volume was calculated as 0.593 ± 0.022 *μ*l using three independent, freshly prepared MSCs. These results confirmed the microfluidic length scale and approximate volume available for liquid flow inside MSCs. It was noteworthy that careful sample preparation for electron miicroscopy turned out to be important in order to obtain more precise microchannel section cuts (both 45^*°*^and 90^*°*^angles) (Figure 7); and to view surface morphology pertaining to inner sections of the MSCs (Figure 8) at necessary mode of magnification (low magnification (LM) or normal (SEM) mode).

**Figure 6:**
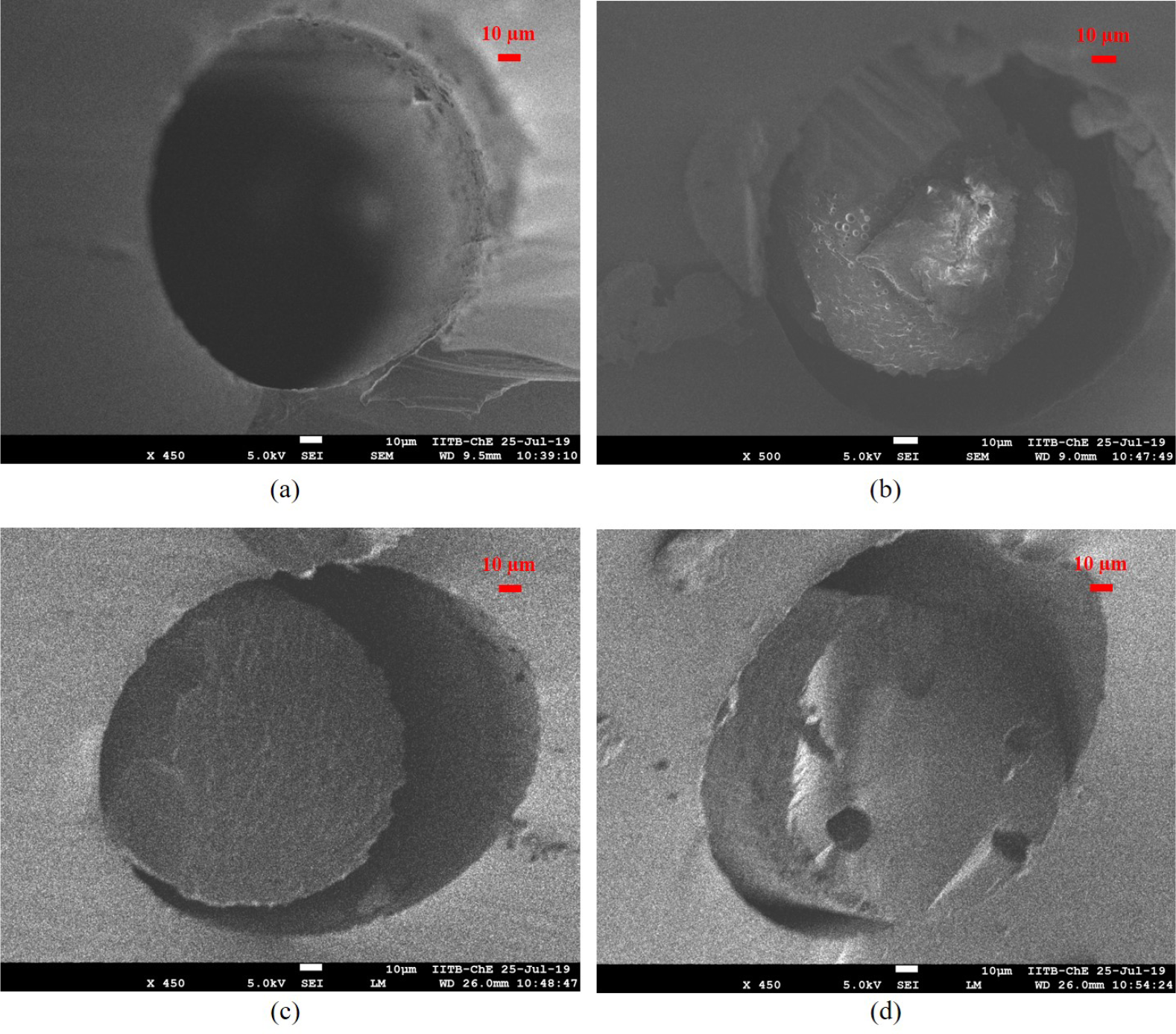
Scanning electron micrographs of channel cross-section for different acrylic copolymer concentrations (in (w/v)%) (450/500X magnification, 45 ° section cut): (a) untreated microchannel (b) MSC (5%) (c) MSC (10%) (d) MSC (15%). Each of these SEM images was obtained using independent microchannels. The scale bar represents 10 *μ*m in all panels.

**Figure 7:**
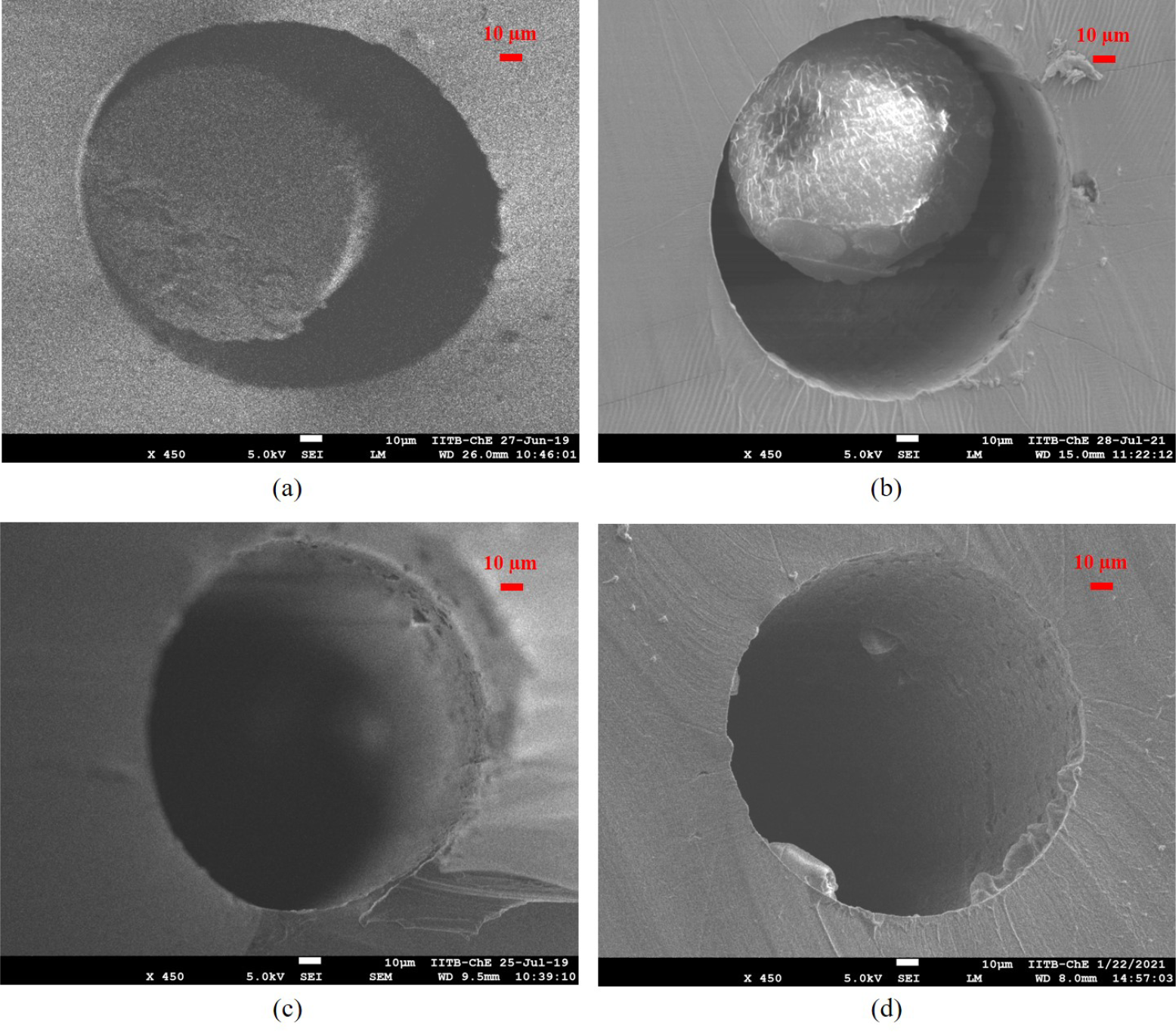
Scanning electron micrographs of MSC cross-section for 10% acrylic copolymer concentration (in (w/v)%) (450X magnification): (a) 45 ° section cut (b) 90 ° section cut. For comparison, micrographs of untreated microchannels ((c) and (d)) at corresponding section cut are provided. Each of these SEM images was obtained using independent microchannels. The scale bar represents 10 *μ*m in all panels.

**Figure 8:**
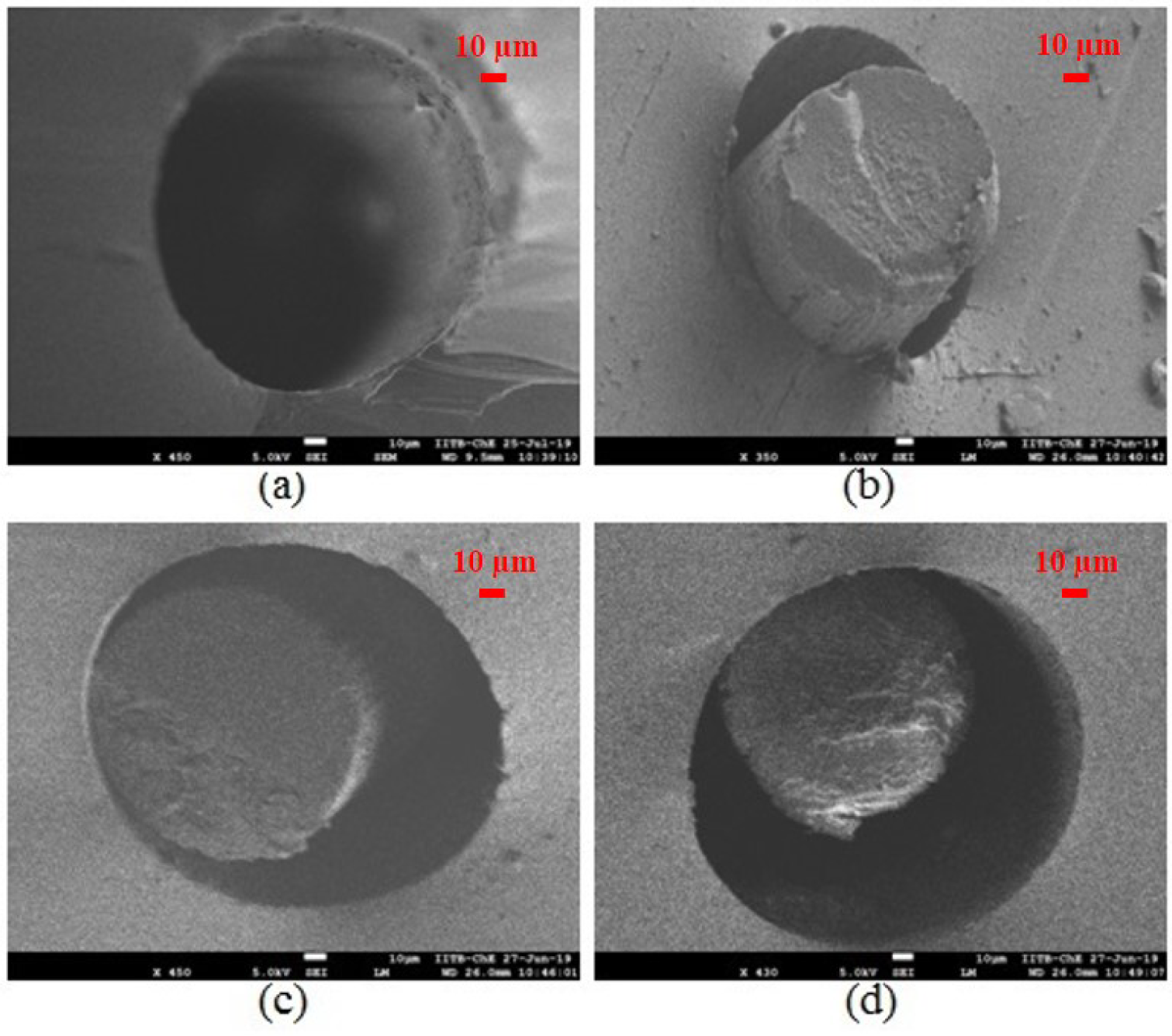
Scanning electron micrographs of column cross-section for 10% acrylic copolymer concentration (in (w/v)%) at different distance from the column inlet (350/450X magnification, 45 ° section cut): (a) untreated microchannel (20 mm away) (b) MSC (10 mm) (c) MSC (20 mm) (d) MSC (30 mm). Each of these SEM images was obtained using independent microchannels. The scale bar represents 10 *μ*m in all panels.

**Figure 9:**
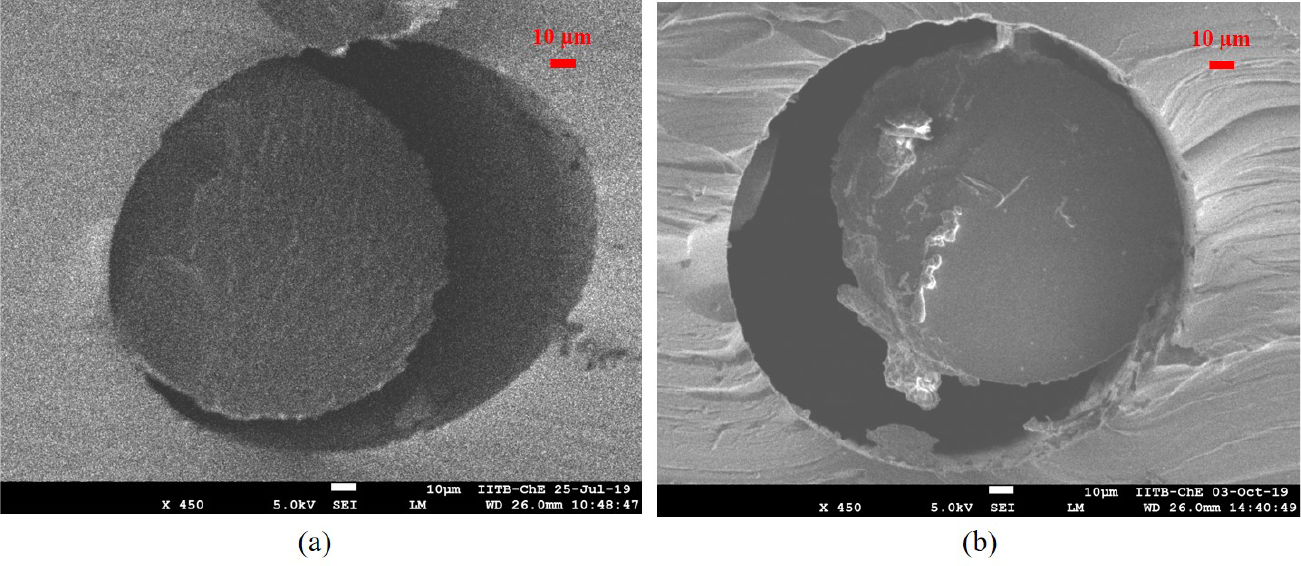
Scanning electron micrographs of MSC cross-section for 10% acrylic copolymer concentration (in (w/v)%) (a) on treatment with phosphate buffer (450X magnification, 45 ° section cut): untreated (no treatment with phosphate buffer) (b) 20 mM phosphate buffer, pH 2.8 (strongly acidic condition). Each of these SEM images was obtained using independent microchannels. The scale bar represents 10 *μ*m in all panels.

### 3.2. Channel visualization

From electron microscopy and preparation of samples with 45^*°*^section cut, the internal details of stationary phase (or monolith) formed by ACP inside the MSCs were further analyzed. As stated in the earlier section, the annular diameter (*D*_c_ − *D*_m_) was around 70 *μ*m in the case of 10% (w/v) ACP concentration, where *D*_c_ ∼ 170 ± 2 *μ*m (measured from 100 different microchannels (Leica DM500 and LAS 3.4 Application software)) and *D*_m_ ∼ 100 *μ*m (measured from SEM images (ImageJ, NIH, USA)). With varying ACP concentration from 5 to 15% (w/v), it was interesting to observe that micropore size also varied from large pores to smaller pores (Figure 10 and image analysis of the SEM images wherever possible). It was consistent with the observations on a microporous region recently reported with ACP surface (or ACP-modified PDMS surface) by our research group [26].

**Figure 10:**
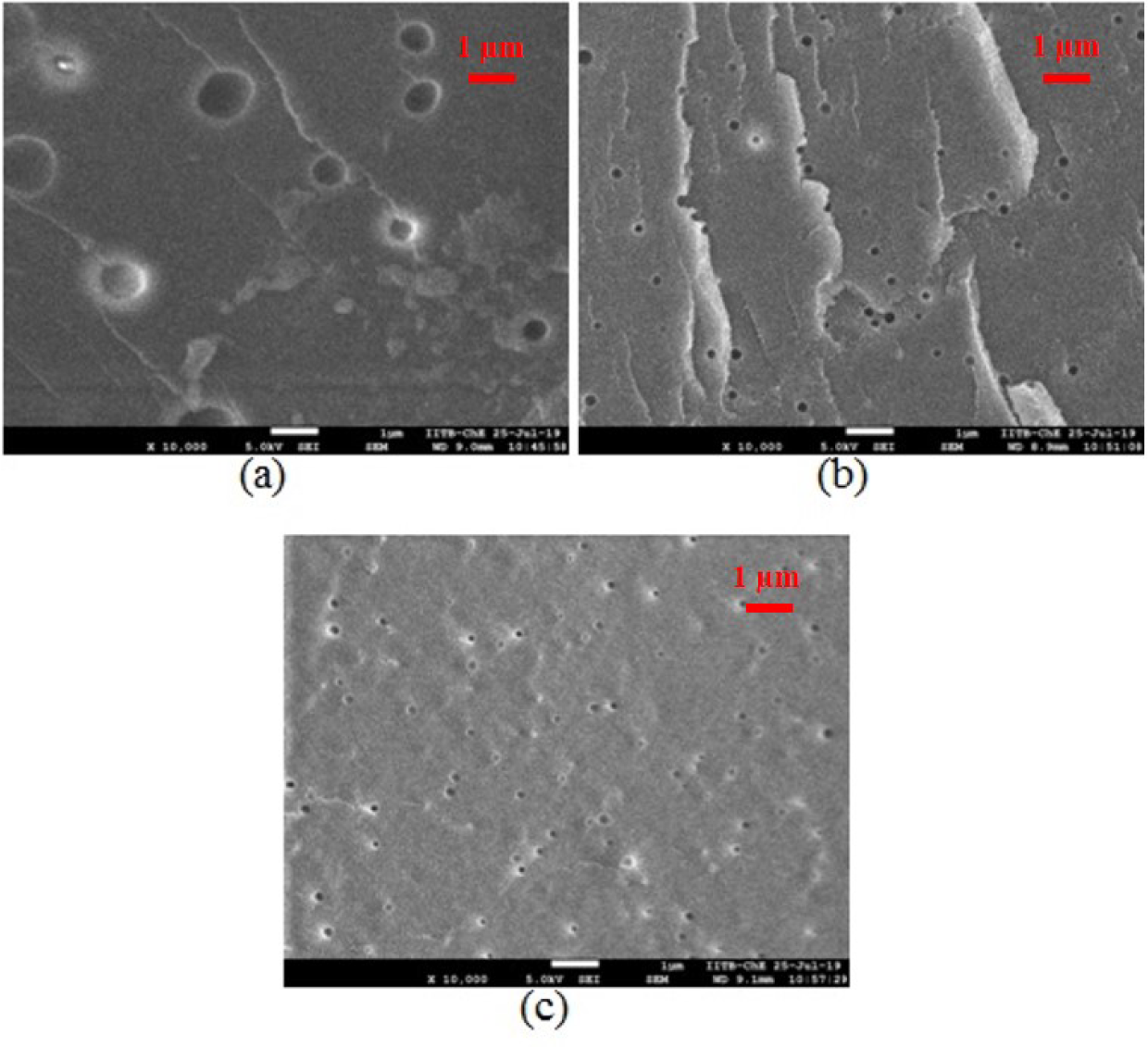
Scanning electron micrographs of micropores on monolith bed inside moon-shaped channel (MSC) after monolith formation with different acrylic copolymer concentrations (in %(w/v)) (10000X magnification, 45 ° section cut): (a) 5% (b) 10% (c) 15%. Each of these SEM images was obtained using independent microchannels. The scale bar represents 1 *μ*m in all panels.

To visualize analyte flow inside a freshly prepared MSC, colored tracer solution namely, blue dextran was used to depict the channel void volume available. The blue-colored region visualized from a mounted camera (Leica DM500) (as top view of MSC) (Figure 11) indicated liquid flow inside MSC. The above results, in combination with likely estimates of size of micropores inside MSC; have been summarized to describe the surface morphology of the functional surface (ACP in the form of monolithic stationary phase) and flow visualization inside MSCs.

**Figure 11:**
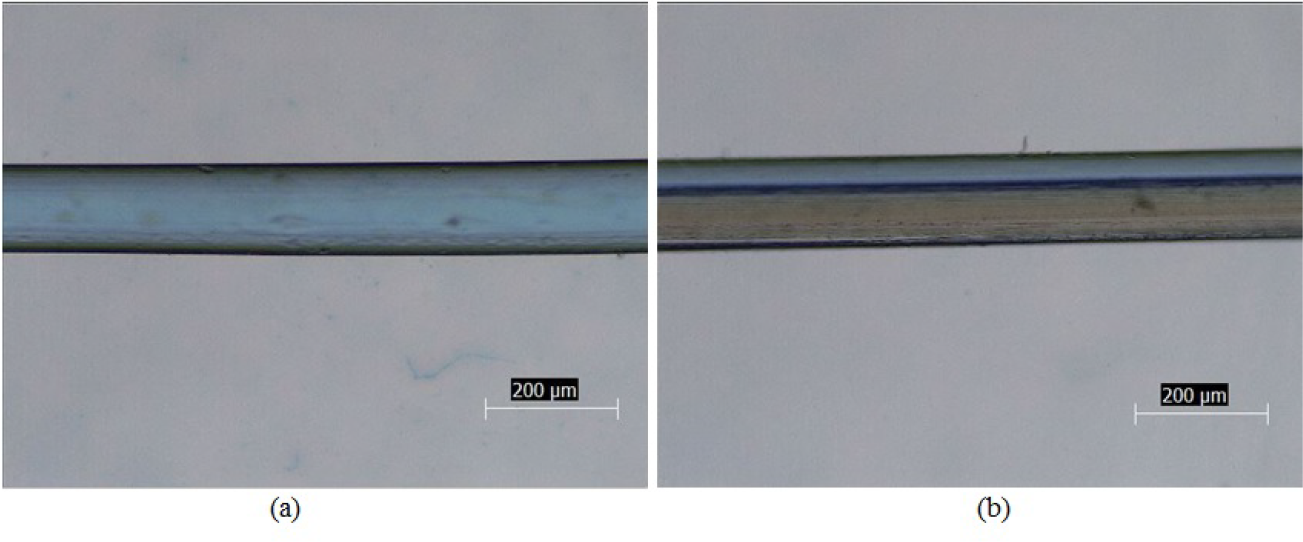
Optical micrographs showing mid-section of moon-shaped channel (MSC) for 10% acrylic copolymer concentration (in (w/v)%) along its length (a) before and (b) after monolith formation; using flow of tracer (35.37 mg ml−1 blue dextran) (10X magnification). The scale bar represents 200 *μ*m in all panels.

### 3.3 Estimate of void volume in MSCs

With step input of tracer as analyte, elution profiles or F-curves are obtained from breakthrough experiments through online monitoring system. For estimating void volume of MSC, a concentration step input was considered and breakthrough point (typically at 10% tail height) noted down from S-shaped curve (as shown in Figure 12). The expression for above estimate was given by Equation 5.

**Figure 12:**
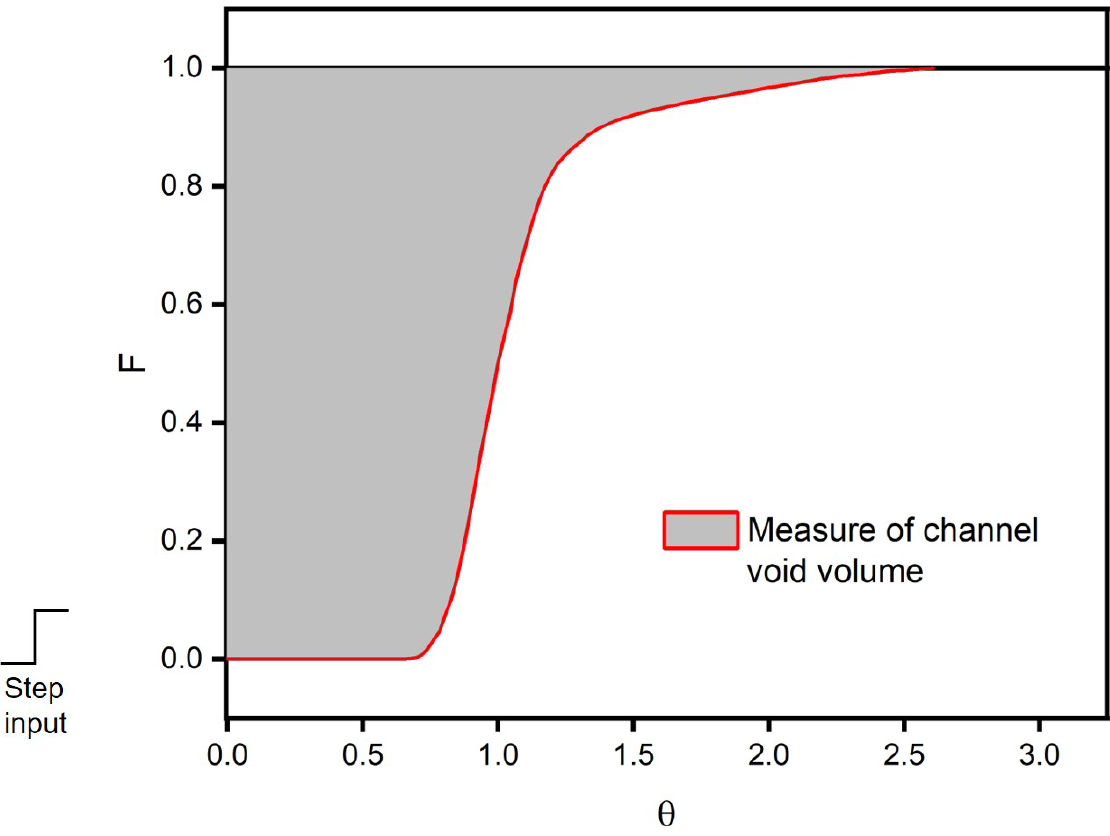
Tracer breakthrough curve for measure of void volume (shaded area) in moon-shaped channel (MSC). Feed: blue dextran (2000 *μ*g ml^−1^) in buffer MB (20 mM phosphate buffer, pH 6.4), stationary phase: 10%(w/v) acrylic copolymer (ACP), eluent: buffer MB (20 mM phosphate buffer, pH 6.4), inlet flow rate: 5 *μ*l min^−1^, analysis time: 2 hours (n=3, performed with independent MSCs).

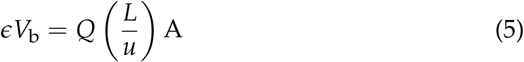

where *ϵV*_b_ void volume of MSC (*μ*l), *Q* - inlet mobile phase flow rate (*μ*l min^−1^), *L* - channel length (mm), *u* - superficial flow velocity (mm min^−1^) and A - dimensionless shaded area from tracer breakthrough curve (refer Figure 12).

Whereas, the expression for geometrical calculations was given by:

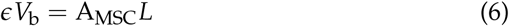

where flow area of cross-section (mm^2^), 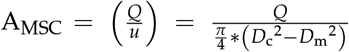 - channel diameter (mm), *D*_m_ - monolith diameter (mm), *L* - channel length (mm)

Note that the shaded area, A provides estimate of void volume only and thus, it does not include volume of porous region inside MSC (or porous region of ACP as shown in Figure 10 is not included with void volume here).

Based on the above estimate from the tracer breakthrough curve experimentally (Equation 5), column void volume for a given MSC was nearly 0.639 ± 0.014 *μ*l. It corresponds to similar value obtained from geometrical calculations (Equation 6) and hence validated with an independent set of at least three MSCs. Since laminar flow inside MSCs and its flow length (40 mm) were known, an estimated pressure drop of ∼ 2 bar indicated the low pressure operating conditions that prevailed during the tracer breakthrough analysis.

Studies with blue dextran (tracer) were performed to validate each MSC fabricated with corresponding residence time distribution (RTD) and to demonstrate contribution of mass diffusion towards peak tailing (Figure 13). A wider RTD was found to be associated with blue dextran due to slow mass diffusion. With step input of tracer as analyte, elution profiles or F-curves are obtained from breakthrough experiments through online monitoring system with sharp peak front. Considering that blue dextran had high molecular weight and high diffusion coefficient, delay in elution was observed along with potential dispersion characteristics in the given microfluidic system when compared to the case of system volume only (Figure 14). These studies indicated that the contribution of MSCs to dispersion inside the system was relatively lower than that from the system volume similar to that of other protein chromatography systems [28] available commercially.

**Figure 13:**
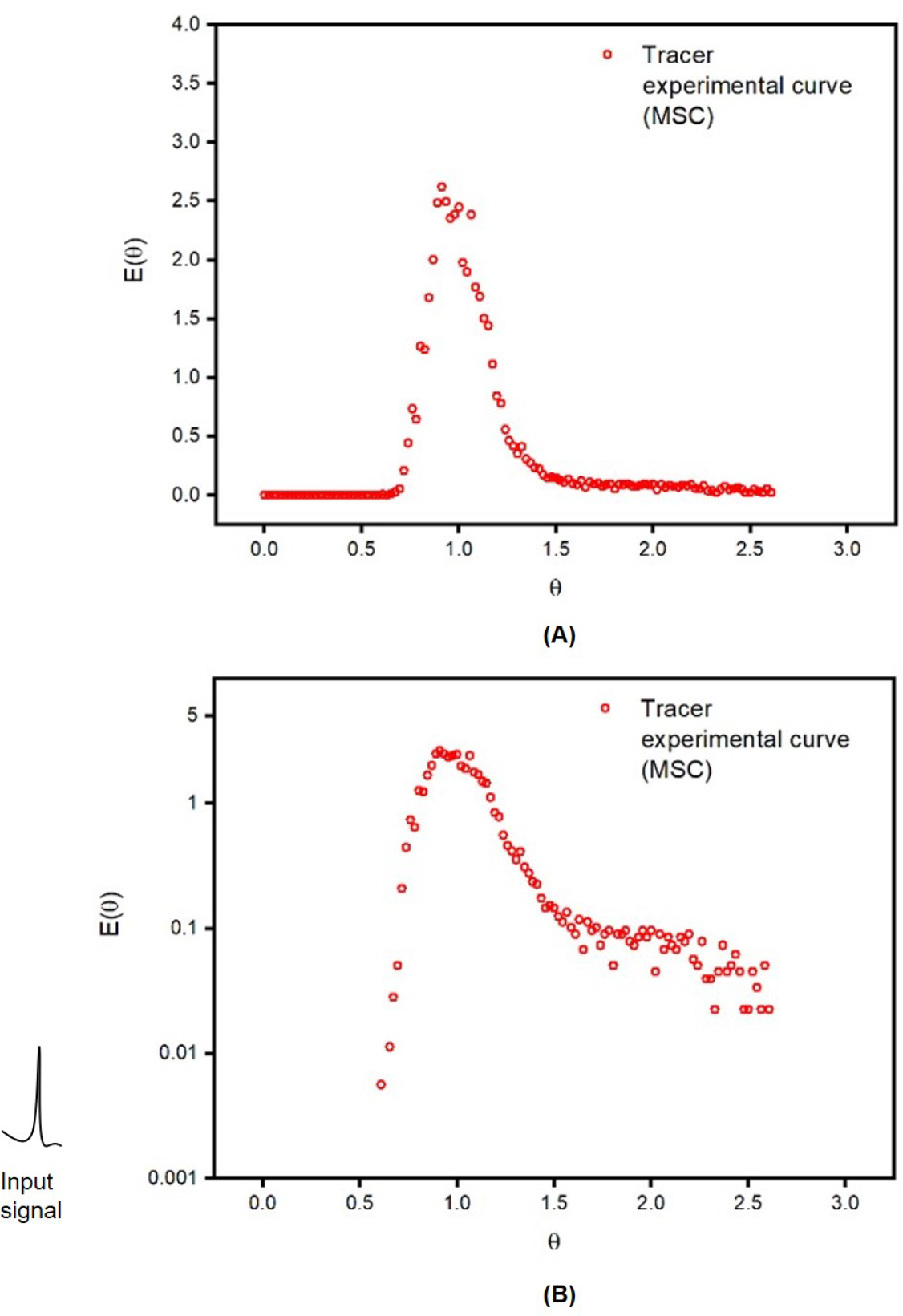
Tracer residence time distribution (RTD) for moon-shaped channel (MSC) (a) E-curve semi-logarithmic plot of E-curve (for zoom-in view of peak tail). Feed: blue dextran (2000 *μ*g ml^−1^) in buffer MB (20 mM phosphate buffer, pH 6.4), stationary phase: 10%(w/v) acrylic copolymer (ACP), eluent: buffer MB (20 mM phosphate buffer, pH 6.4), inlet flow rate: 5 *μ*l min^−1^, analysis time: 2 hours (n=5, performed with independent MSCs).

**Figure 14:**
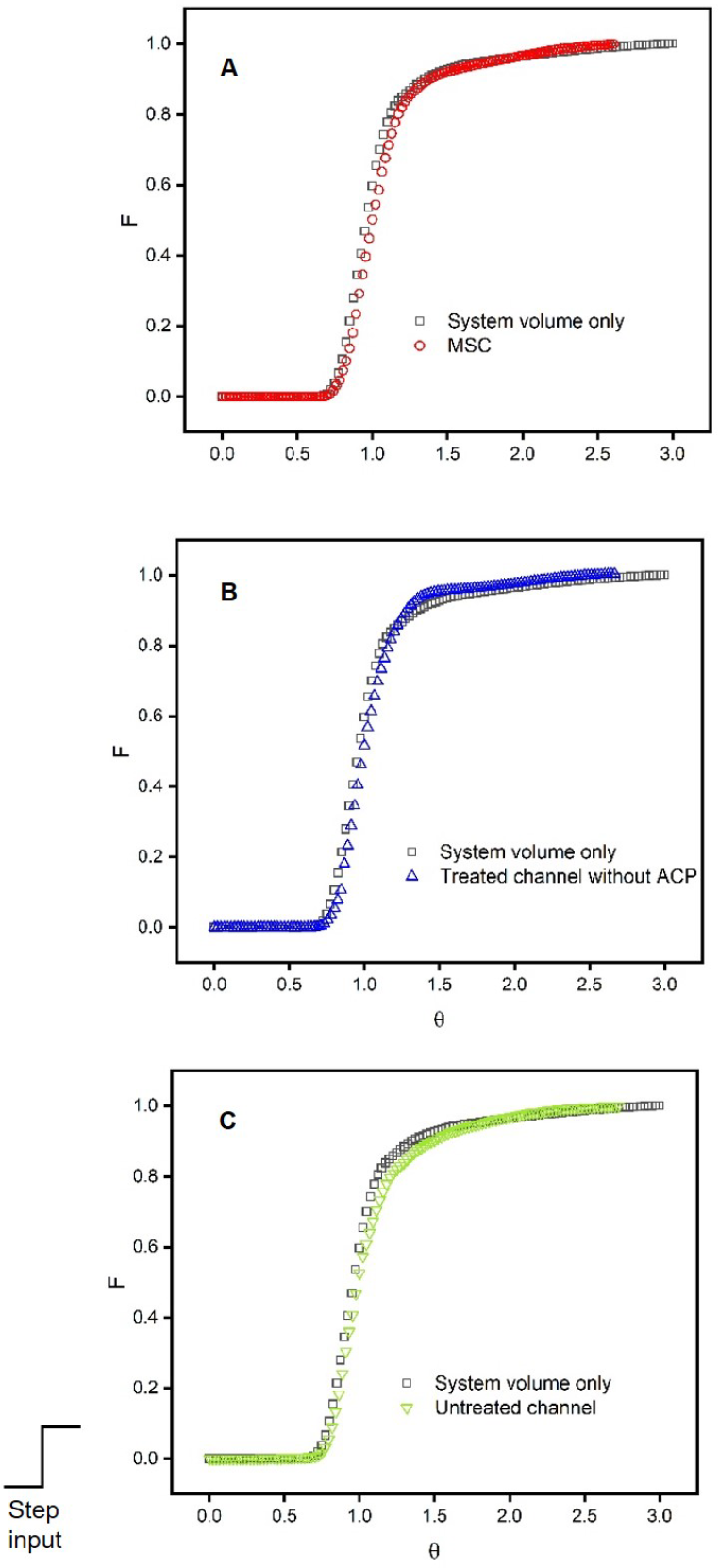
Comparison of tracer breakthrough curves with connecting microchannel (microchannel connected inline with system volume) or without connecting microchannel (system volume only). The type of microchannel connected in each case is as follows: (A) moon-shaped channel (MSC) (n=5), (B) treated channel without adsorbent (or ACP) (n=4) (C) untreated channel (or empty channel) (n=5). Feed: blue dextran (2000 *μ*g ml^−1^) in buffer MB (20 mM phosphate buffer, pH 6.4), stationary phase: 10%(w/v) acrylic copolymer, eluent: buffer MB (20 mM phosphate buffer, pH 6.4), inlet flow rate: 5 *μ*l min^−1^, analysis time: 2 hours (each trial performed with independent microchannels).

### 3.4. Protein breakthrough analysis

With step input of human hemoglobin as model protein, elution profiles or F-curves are obtained from breakthrough experiments through online monitoring system. Peak tailing after nearly 70% of total protein feed clearly indicated that dispersion previously observed with tracer was captured during the protein breakthrough analysis (Figure 15). Based on the above estimate from the protein breakthrough curve experimentally (Equation 5), column void volume for a given MSC was nearly 0.662 ± 0.065 *μ*l (Table 2). It correspond to similar values obtained from both geometrical calculations (Equation 6) and tracer breakthrough analysis. All of the above results were also validated with an independent set of at least three MSCs.

**Table 2:** Estimates of void volume in moon-shaped channel.

| Source of estimate | Channel void volume* ($\mu\text{l}$) |
| --- | --- |
| Geometry | $0.593 \pm 0.022$ |
| Experiments with blue dextran | $0.639 \pm 0.014$ |
| Experiments with human hemoglobin | $0.662 \pm 0.065$ |

\*Estimated from three independent, freshly prepared moon-shaped channels (MSCs)

**Figure 15:**
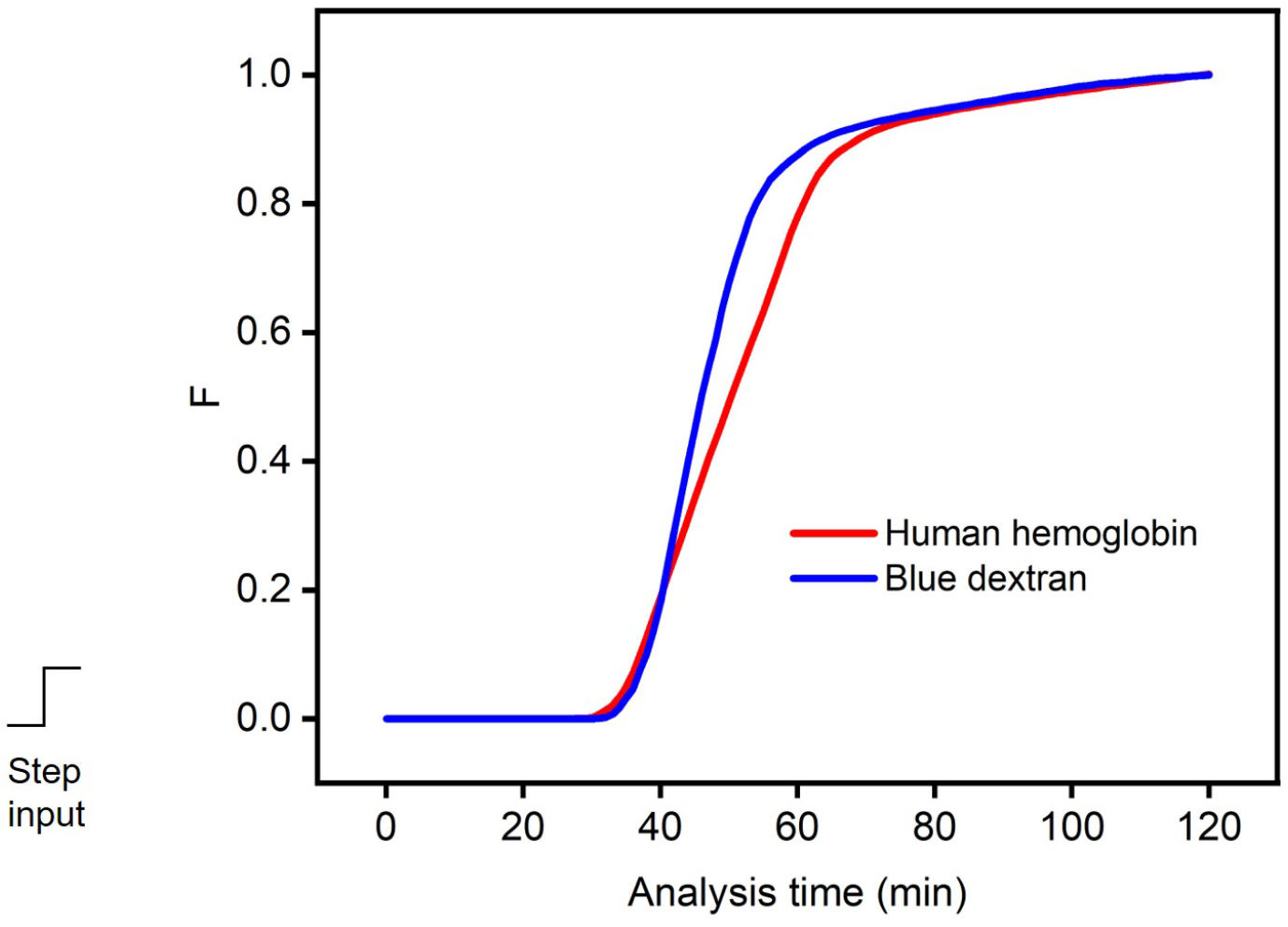
Experimental protein breakthrough curve in moon-shaped channel (MSC). Tracer breakthrough curve is superimposed onto the protein breakthrough curve for comparison of tracer residence time and retention time of protein fractions. Feed: human hemoglobin (1000 *μ*g ml^−1^) in phosphate buffer, stationary phase: 10%(w/v) acrylic copolymer (ACP), presaturation buffer: buffer AB (20 mM phosphate buffer, pH 9.3), mobile phase buffer for elution: buffer MB (20 mM phosphate buffer, pH 6.4), inlet flow rate: 5 *μ*l min^−1^, analysis time: 2 hours (averaged from n=5 trials).

At single pH condition in the acidic range, elution profiles were analyzed using human hemoglobin A_0_ (or HbA_0_) as single protein. A slight delay in retention time was found relative to residence time of the microfluidic system. Tracer RTDs were compared to the protein elution profiles (acidic pH 2.5 only) and the above delay was validated (Figure 16). This turned out to be a favorable condition for hemoglobin chromatography and hence, more potential trials in presence of different hemgolobin variants were required to demonstrate case studies with complex protein mixtures some of which are underway in our research group.

**Figure 16:**
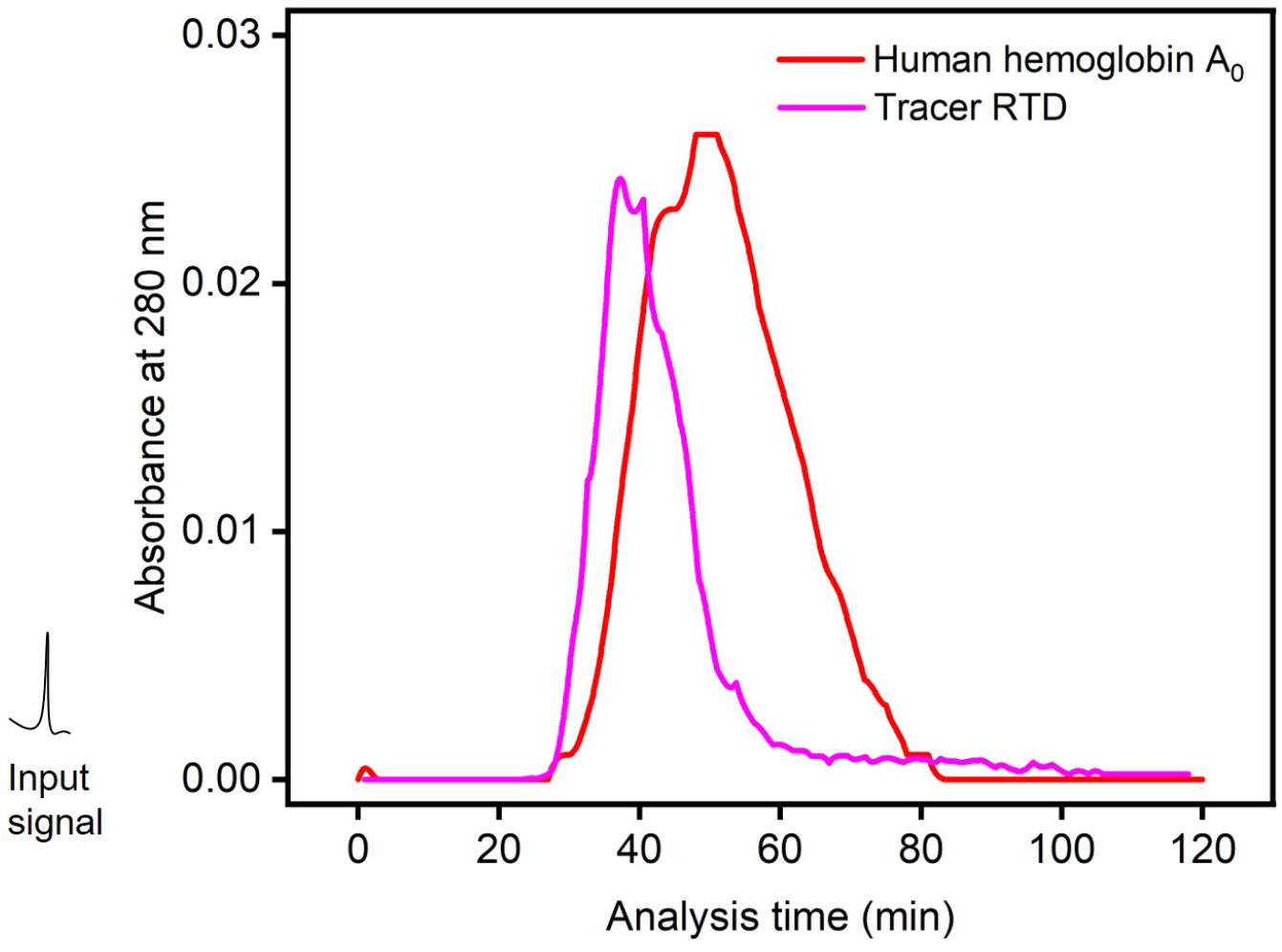
Experimental elution profiles with single protein in moon-shaped channel (MSC) at single pH condition (acidic pH of 2.5). Tracer residence time distribution (RTD) is superimposed onto the elution profiles for comparison of tracer residence time and retention time of protein fractions. Feed: human hemoglobin A_0_ (or HbA_0_) (1.7 *μ*g total protein in 20 mM phosphate buffer, pH 6.4), stationary phase: 10%(w/v) acrylic copolymer (ACP), presaturation buffer: buffer PB (20 mM phosphate buffer, pH 2.5), mobile phase buffer for elution: buffer MB (20 mM phosphate buffer, pH 2.5), inlet flow rate: 5 *μ*l min^−1^, analysis time: 2 hours (averaged from triplicate independent experiments with freshly prepared MSCs).

### 3.5. Reconditioning of MSCs

Moon-shaped channels (MSCs) were fabricated using acrylic copolymer (ACP) as the monolith material. It led to formation of stationary phase inside the channels that facilitated its interactions with proteins. In order to demonstrate reconditioning, fabrication procedure was revisited and the utility of organic solvent in dissolution of ACP thoroughly studied. It turned out that use of organic solvent after MSC fabrication affected the monolith significantly.

Acrylic copolymer inside MSC was found to be washed away after treatment with excess of organic solvent (Figure 17). Since the monolith is made up of acrylic copolymer that has high solubility in tetrahydrofuran (organic solvent), it gets redissolved and the channel could be possibly regenerated with fresh porous monolith formation.

**Figure 17:**
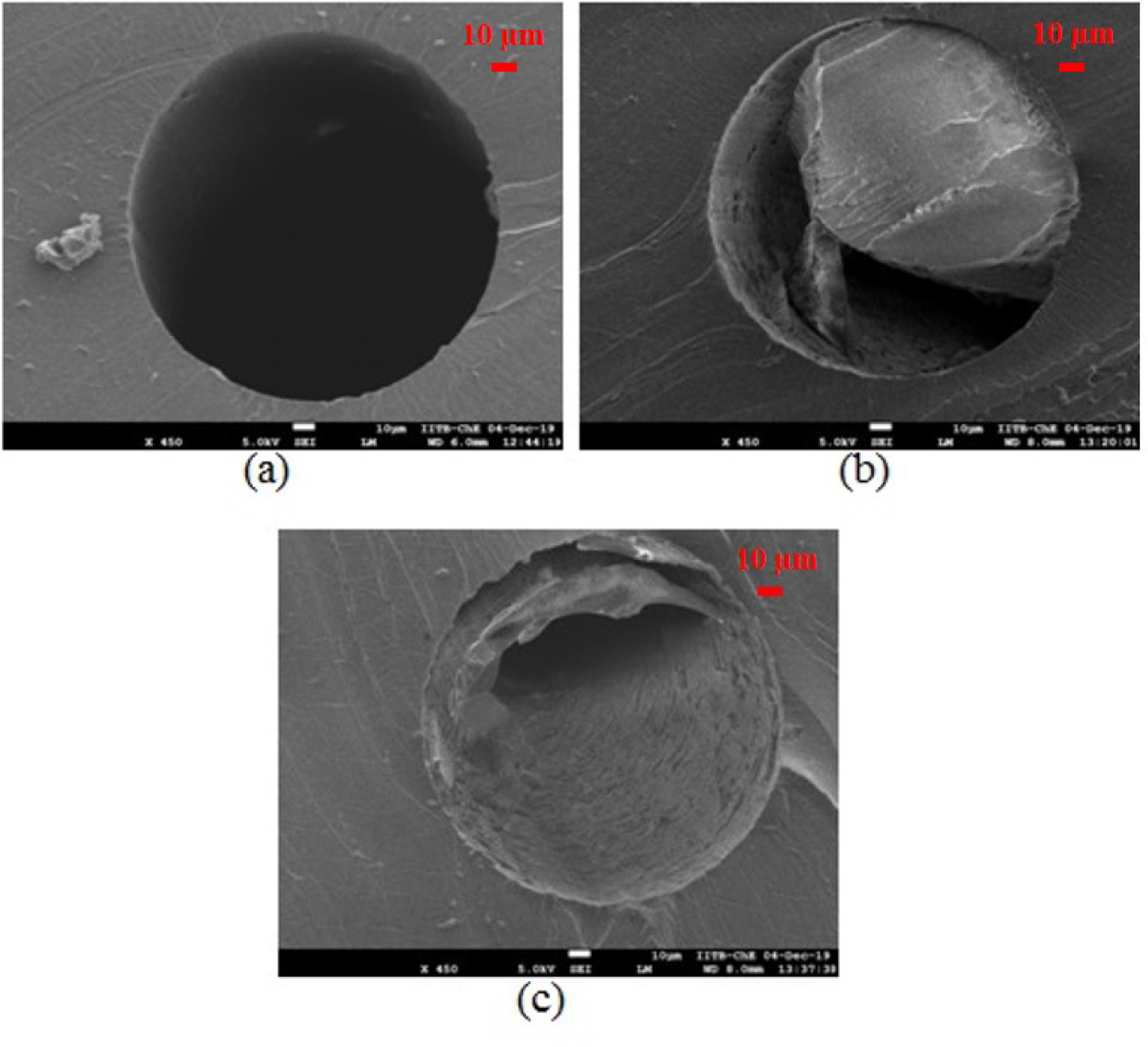
Scanning electron micrographs of column cross-section using acrylic copolymer concentration of 10% (w/v) (450X magnification, 90^*o*^ section cut): (a) untreated microchannel (b) MSC with no treatment using excessive organic solvent (or tetrahydrofuran (THF)) (c) MSC on treatment with excessive THF. Each of these SEM images was obtained using independent microchannels. The scale bar represents 10 *μ*m in all panels.

## 4. Conclusion

A facile microchannel fabrication technique using a combination of templatebased micromolding and formation of stationary phase to obtain chip-based moon-shaped channel (MSC) has been presented in the present work. Polymer monoliths as stationary phase inside the channels are obtained through rapid prototyping and the channels do not require any bonding step. Fine tuning of monolith diameter of moon-shaped channels (MSCs) is possible with change in acrylic copolymer (ACP) concentration. By establishing a microfluidic system with online monitoring, an estimate of channel void volume from breakthrough curves has been experimentally determined using tracer elution profiles. Protein breakthrough analysis indicated the elution profiles in the laminar flow regime as similar to tracer and possible retention of hemoglobin as model proteins. The above case study with single protein could be potentially explored in complex protein mixtures with two or more types of protein to demonstrate microfluidic liquid chromatography at low column pressure drop.

## 6. CRediT author statement

Raghu K. Moorthy: Conceptualization, methodology, validation, formal analysis, investigation, data curation, writing - original draft, visualization. Serena D’Souza: Formal analysis, writing - review & editing, visualization. P. Sunthar: Methodology, formal analysis, resources, data curation, writing - review & editing, visualization, supervision, project administration. Santosh B. Noronha: Methodology, formal analysis, resources, writing - review & editing, visualization, supervision, project administration, funding acquisition.

## 7. Acknowledgements

We are thankful to the instrument facilities provided by Machine Lab (Tata Centre for Technology and Design), Continuous Flow Chemistry Lab (Department of Chemistry), Material Characterization and Testing Lab (Department of Chemical Engineering), Sophisticated Analytical Instrumentation Facility (SAIF) and Industrial Research and Consultancy Centre (IRCC) at IIT Bombay. A special mention about Ms. Meena Menghrajani (formerly Research Engineer, Department of Chemical Engineering/IRCC), who assisted us with her expertise in field emission gun-scanning electron microscopy (FEG-SEM) during sample preparation of cylindrical microchannels. One of the co-authors (RKM) duly acknowledges financial support received from Department of Science and Technology (DST), Government of India.

